# MarpolBase Expression: A Web-based, Comprehensive Platform for Visualization and Analysis of Transcriptomes in the Liverwort *Marchantia polymorpha*

**DOI:** 10.1101/2022.06.03.494633

**Authors:** Shogo Kawamura, Facundo Romani, Masaru Yagura, Takako Mochizuki, Mika Sakamoto, Shohei Yamaoka, Ryuichi Nishihama, Yasukazu Nakamura, Katsuyuki T. Yamato, John L. Bowman, Takayuki Kohchi, Yasuhiro Tanizawa

## Abstract

The liverwort *Marchantia polymorpha* is equipped with a wide range of molecular and genetic tools and resources that have led to its wide use to explore the evo-devo aspects of land plants. Although its diverse transcriptome data are rapidly accumulating, there is no extensive yet user-friendly tool to exploit such a compilation of data and to summarize results with the latest annotations. Here, we have developed a web-based suite of tools, MarpolBase Expression (MBEX, https://marchantia.info/mbex/), where users can visualize gene expression profiles, identify differentially expressed genes, perform co-expression and functional enrichment analyses, and summarize their comprehensive output in various portable formats. Using oil body biogenesis as an example, we demonstrated that the results generated by MBEX were consistent with the published experimental evidence and also revealed a novel transcriptional network in this process. MBEX should facilitate the exploration and discovery of the genetic and functional networks behind various biological processes in *M. polymorpha,* and promote our understanding of the evolution of land plants.

## Introduction

The liverwort *Marchantia polymorpha* has become one of the most actively studied model plants in recent years. *M. polymorpha* is dioicous, has a haploid gametophyte-dominant life cycle, and has two modes of reproduction: asexual reproduction through clonal propagules called gemmae, and sexual reproduction through sperm and eggs generated in reproductive organs, antheridia and archegonia, respectively. Liverworts are one of the three extant groups of Bryophyta, which forms a sister group to the other lineage of Embryophyta, the vascular plants. Therefore, comparisons between *M. polymorpha* and the other lineages provide insights into evo-devo aspects of land plants.

The genome of *M. polymorpha* subsp. *ruderalis* has been sequenced (Bowman et al., 2017; Montgomery et al., 2020)) and assembled into chromosomes (Montgomery et al., 2020). Within the *M. polymorpha* genome, there exists less redundancy among regulatory genes while most transcription factor families are retained, suggesting that many aspects of *M. polymorpha* share with other land plant species a common regulatory toolkit that is representative of the core elements present in the earliest embryophytes. Studies using *M. polymorpha* as a model have provided insights into the conservation and diversity of responses and regulatory mechanisms in land plant evolution and their evolutionary origins (Kohchi et al. 2021).

In the past decade, a wide range of molecular and genetic tools and resources for *M. polymorpha* have been developed, including efficient *Agrobacterium-mediated* transformation, a series of convenient vectors, and efficient genome editing using the CRISPR/Cas9 system (reviewed by Kohchi et al., 2021). A website dedicated to *M. polymorpha,* MarpolBase (https://marchantia.info), has been developed and is continually updated for the research community (Bowman et al., 2017). MarpolBase provides a comprehensive set of data resources, such as sequence data, gene models, and annotations, including KEGG, KOG, Pfam, and GO terms, serving as a data hub for genome-based studies, especially for those utilizing next-generation sequencing (NGS) techniques. MarpolBase also serves as a repository for *M. polymorpha* gene names to maintain consistency and avoid redundancy and confusion in the scientific literature.

The rapid spread of *M. polymorpha* as a model system allowed numerous research advances, many employing gene expression analyses (Kohchi et al., 2021; Bowman, 2022). It is noteworthy that RNA-seq has become a routine lab procedure to capture global gene expression patterns due to its simplicity and cost-effectiveness, and it is indeed accelerating studies using *M. polymorpha.* RNA-seq enables the determination of spatio-temporal and quantitative expression of genes of interest under different environmental conditions, in specific tissues, organs, or developmental stages, or in distinct genotypes, providing valuable clues to elucidate gene function.

An ever-growing number of RNA-seq datasets have been accumulating in the Sequence Read Archive (SRA) maintained by the International Nucleotide Sequence Database Collaboration (INSDC) (Katz et al., 2022). Since no restrictions are placed on the reuse and redistribution of the data archived in the INSDC, there are many ‘secondary databases’, where sequence data from the SRA are reanalyzed and processed to better understand the entirety of public data, thereby providing novel insights. The accumulation of RNA-seq data from a wide range of conditions, tissues, and mutants enables us to utilize the data for co-expression analyses to predict a particular set of genes that act coordinately, and also to generate hypotheses about gene functions. Co-expression analysis, which assumes that genes with relevant functions are expressed in a similar spatial and temporal pattern even under different conditions, has indeed revealed many regulatory relationships. User-friendly platforms enable easy community access to a large array of data for co-expression analyses. Successful examples are the Arabidopsis RNA-seq database (ARS), which serves as a platform for comprehensive expression analysis generated from intensive reanalysis of more than 20,000 Arabidopsis datasets from public resources; ATTED-II (Obayashi et al., 2018), another co-expression database for Arabidopsis as well as some crop plants; and COXPRESdb (Obayashi et al., 2019), which provides co-regulated gene networks in yeasts and animals. These provide a web-based interface that enables easy browsing, analysis, and visualization. Also available are those for other model plants, such as *Physcomitrium patens* (Perroud et al., 2018) and *Selaginella moellendorffii* (Ferrari et al., 2020). These databases created by the reuse of public sequencing data should accelerate biological studies on these species.

To take advantage of accumulating RNA-seq data derived from *M. polymorpha,* we have collected and analyzed the existing datasets and developed an interactive database, named MarpolBase Expression (MBEX, https://marchantia.info/mbex/), with a user-friendly interface that allows comprehensive analysis and web-based visualization of public RNA-seq data. This database provides tools that enable users to visualize gene expression levels in various portable formats, analyze co-expression data, including co-expression networks, identify differentially expressed genes, and perform functional enrichment analysis. Recently, similar expression and co-expression databases have been developed for *M. polymorpha*, emphasizing abiotic stress responses and diurnal gene expression, as well as organogenesis and reproduction gene expression profiles (Julca et al., 2021; Tan et al., 2021). They are provided as part of the ‘electronic Fluorescence Pictographs’ (eFP) browser (http://bar.utoronto.ca) and the Evorepro database (https://evorepro.sbs.ntu.edu.sg). While MBEX has some overlap with these databases, it provides an all-in-one analysis platform closely linked with MarpolBase, and thus continuously incorporates annotation updates as well as additional RNAseq datasets as they are deposited in NCBI and other sequence archives. As demonstrated in a later section, users can complete the entire expression analysis on a single website using the tools provided. Further, gene annotations have been updated to the latest reference genome for *M. polymorpha* (MpTak or ver. 6.1 genome that includes genes on both sex chromosomes [U and V]) by assigning functions and curated names to more than 2,300 genes, which have been made available from MarpolBase. Finally, future updates of gene annotations made for MarpolBase will be imported to MBEX without delay. MBEX should facilitate functional and evolutionary analyses of the genes in *M. polymorpha.*

## Results & Discussion

### Database Construction

To construct an expression database of *M. polymorpha,* we retrieved 460 transcriptome datasets from the SRA. They represent all major tissues and organs covering the entire *M. polymorpha* life cycle; *i.e.,* the vegetative gametophyte (thallus), male and female sexual organs (antheridiophore and archegoniophore, respectively), male and female gametangia (antheridium and archegonium, respectively), sporophytes, spores, asexual reproductive organs (gemma and gemma cup), and apical cells. The collection also includes 117 transcriptomes from mutants and experiments with time courses involving drug treatments and bacterial infection, including the comprehensive collection of RNA-seq data examined previously (Flores-Sandoval et al., 2018). We processed all RNA-seq data with the same pipeline and obtained the Transcripts Per Million (TPM) and raw count matrix. After filtering out samples with low mapping rates, we constructed the co-expression matrix from the TPM matrix. (Liesecke et al., 2018; Obayashi et al., 2018). This database provides visualization tools for gene expression profiles and co-expression networks, as well as analytical tools for NGS data, such as differentially expressed gene (DEG) analysis, functional enrichment analysis, and set relation analysis (Fig. 1A).

**Fig. 1.**
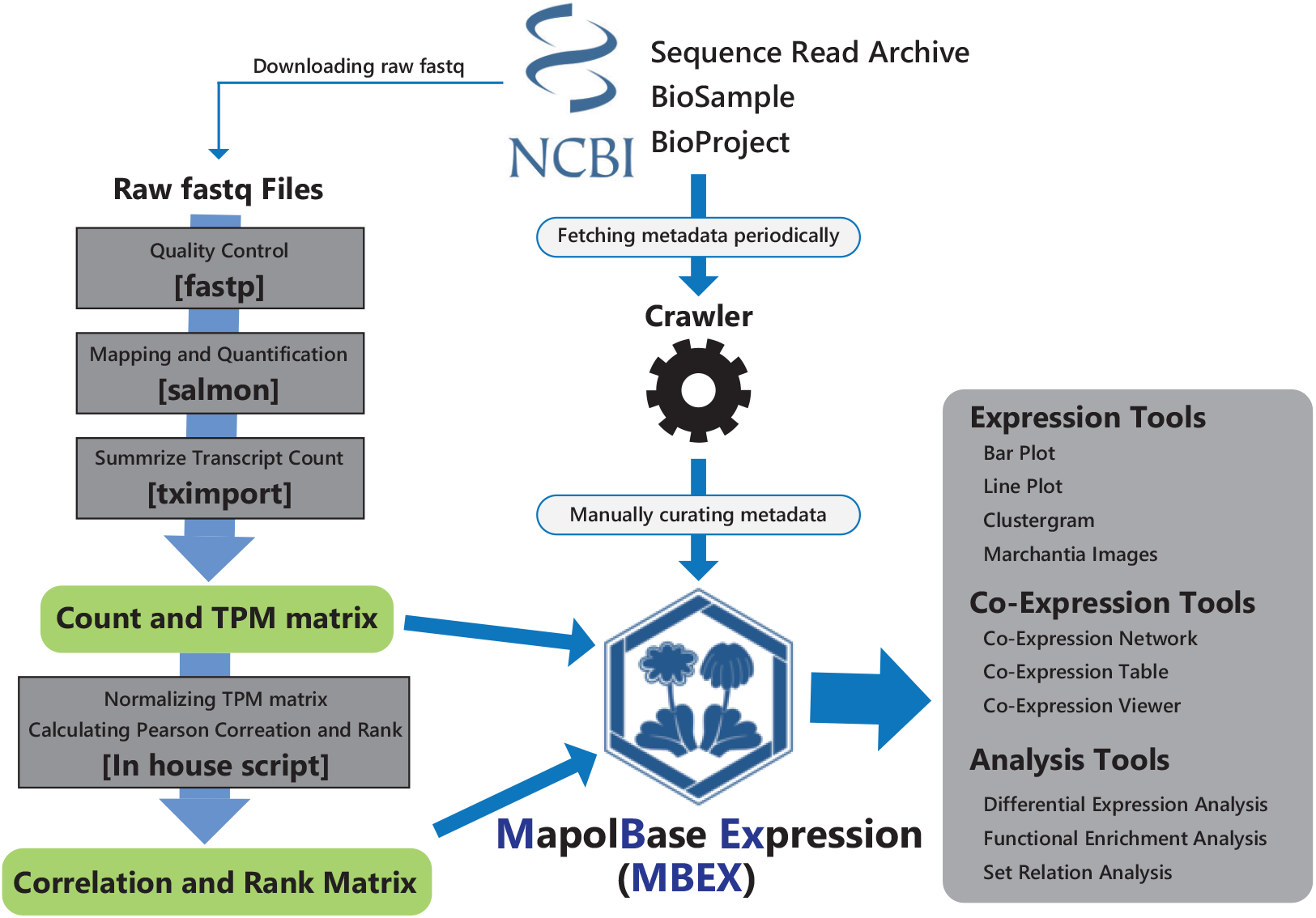
Schematic overview of the construction of MarpolBase Expression (MBEX).

To enhance these main features, we also implemented the additional features ‘OrthoPhyloViewer’, ‘Data Source’, and ‘Co-Expression Viewer’ (Supplementary text). ‘OrthoPhyloViewer’ finds an orthologous group to which a given gene belongs and provides orthologs in a tabular format with a phylogenetic tree to facilitate comparative analysis with other model plant species (Supplementary Fig. S1A, B). ‘Data Source’ helps search for datasets used in MBEX and provides external links to their original data in SRA for download, along with their associated papers (Supplementary Fig. S1C). ‘Co-Expression Viewer’ is a tool to show expression correlations between two genes of interest in all the samples used in MBEX (Supplementary Fig. S1D). It should be noted that this database is kept up to date by periodically acquiring newly deposited RNA-seq data and their metadata through NCBI’s API.

### Improvement of the M. polymorpha genome annotation

The latest version of the *M. polymorpha* genome at MarpolBase (MpTak_v6.1) consists of the autosomal and V chromosomal sequences of the male reference strain Takaragaike-1 (Tak-1) and the U chromosomal sequences from the female reference strain Tak-2 (Iwasaki et al., 2021; Montgomery et al., 2020). Functional annotations for each gene model were also imported to MBEX from MarpolBase. Two-thirds of the genes are annotated with at least one of the Pfam domains, KEGG/KOG orthology, or GO terms (Supplementary Fig. S2). Future updates of gene annotations will be periodically imported from MarpolBase, so the latest annotations can be readily available from MBEX.

From an intensive literature survey, we also manually curated more than 2,300 genes with names, functions, and citations following the guidelines for gene nomenclature established by the community (Bowman et al., 2016). The newly annotated genes were registered to the Marchantia Nomenclature Database (https://marchantia.info/nomenclature/) and made publicly available. In total, there are currently 3,349 manually curated gene annotations, comprising 18.3% of the protein coding genes of the *M. polymorpha* v6.1 genome. In MBEX, users can use most of the tools by querying gene names, e.g., *MpPHY,* as well as gene identifiers (MpGene ID), e.g., Mp2g16090, thereby improving the accessibility and the flexibility of the website.

### Exploratory Data Analysis

We first performed an exploratory analysis to check the validity of the collected RNA-seq data. In hierarchical clustering analysis, most biological replicates were consistently grouped or associated with similar samples (Fig. 2A). The mapping rates of 405 out of 460 samples were over 50%, but 55 samples showed relatively or severely low mapping rates (Supplementary Fig. S3, Supplementary Data). Careful consideration should be taken when using these low mapping rate samples. Further, the principal component analysis showed that the same or similar tissues of *M. polymorpha* had similar transcriptomic profiles (Fig. 2B) regardless of experimental conditions and techniques, such as sequencing method, culture media, and ecotypes, which indicates sufficient reproducibility and consistency of our pipeline. Differences in transcriptomic profiles are more prominent between vegetative and reproductive tissues or gametophytic and sporophytic generations than between different environmental conditions, as shown previously (Flores-Sandoval et al., 2018). Highly divergent transcriptomic profiles between sporophyte and gametophyte were also observed in the *P. patens* transcriptomic atlas (Perroud et al., 2018). Interestingly, samples from early sporelings, developing gemmae, and the apical meristem were grouped, probably due to early sporelings and developing gemmae being rich in meristematic cells. These results also suggest that the apical meristem genetic program differs from that in mature tissues. During reproductive growth, archegonia, antheridia, and sperm show distinct profiles from those of vegetative tissues. Both antheridiophores and archegoniophores clustered together, and consistent with their origin as modified thallus they are located between the clusters of the vegetative gametophyte and gametangia samples (Fig. 2B, Supplementary Fig. S4). Interestingly, the samples from *Mprkd* mutant archegonia (Hisanaga et al., 2021), which do not form mature eggs (Koi et al., 2016; Rövekamp et al., 2016), are more similar to those from gametangiophore tissue than those from normal archegonia, suggesting the egg cell contributes substantially to the wild-type archegonia signature (Fig. 2B, Supplementary Fig. S4). These results are consistent with the fact that the structure of the gametangiophore is similar to that of the thallus (Shimamura, 2016).

**Fig. 2.**
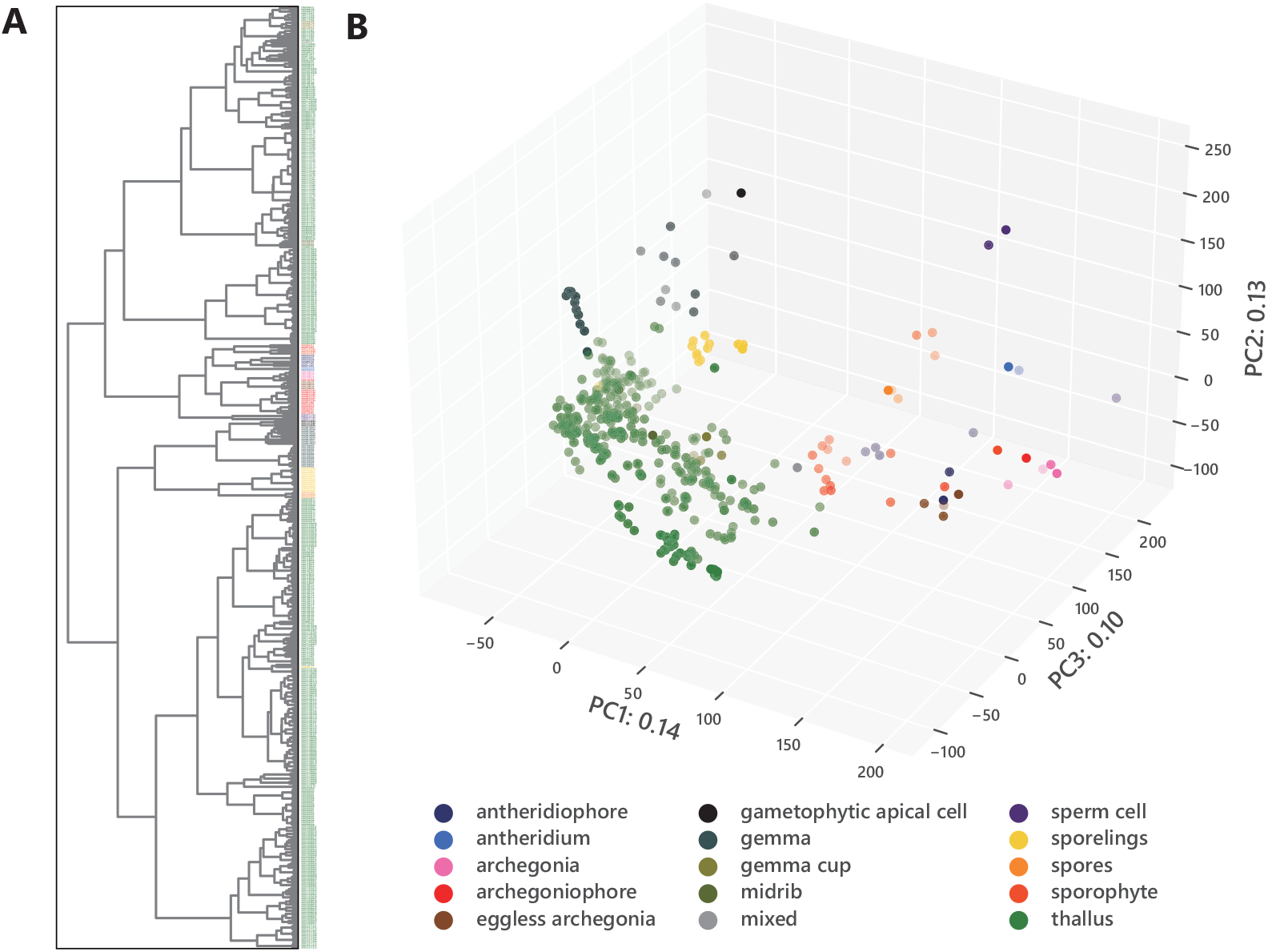
Evaluation of the RNA-seq data used in this study. RNA-seq data collected from SRA were evaluated prior to incorporation into the database. Hierarchical clustering (**A**) and principal component analysis (**B**) of all the 460 samples used in this study. Tissue types are color-coded as shown at the bottom right corner. The number of samples for each tissue is given in Supplementary Table S1.

### Visualization of Gene Expression Profiles

When, where, and under which conditions a gene is expressed can provide insights into gene function. For example, a gene expressed specifically during reproductive growth suggests that the gene is required for a process such as sexual organ formation or germ cell specification. To provide such information, we compiled and visualized the expression levels of each gene in TPM from different samples. In MBEX, expression levels can be visualized as heatmaps represented by images of *M. polymorpha* organs (‘Chromatic Expression Images’), simple plots (‘Bar Plot’ and ‘Line Plot’), or hierarchical clustering images (‘Clustergram’) (Fig. 3A-D). ‘Chromatic Expression Images’ provides an accessible overview of the expression pattern in the entire life cycle of the user-selected gene, similar to the eFP developed for other plant species (Winter et al., 2007). In ‘Bar Plot’, users can visualize the expression levels of a user-selected gene according to user-selected conditions. Similarly, users can select conditions and genes in ‘Line Plot’ and ‘Clustergram’ to view expression patterns. Users have access to detailed information such as the mean and standard deviation with any of these tools. Users can also download the expressionlevel data as images in SVG, PNG, and JPEG formats, and raw TPM values in CSV format.

**Fig. 3.**
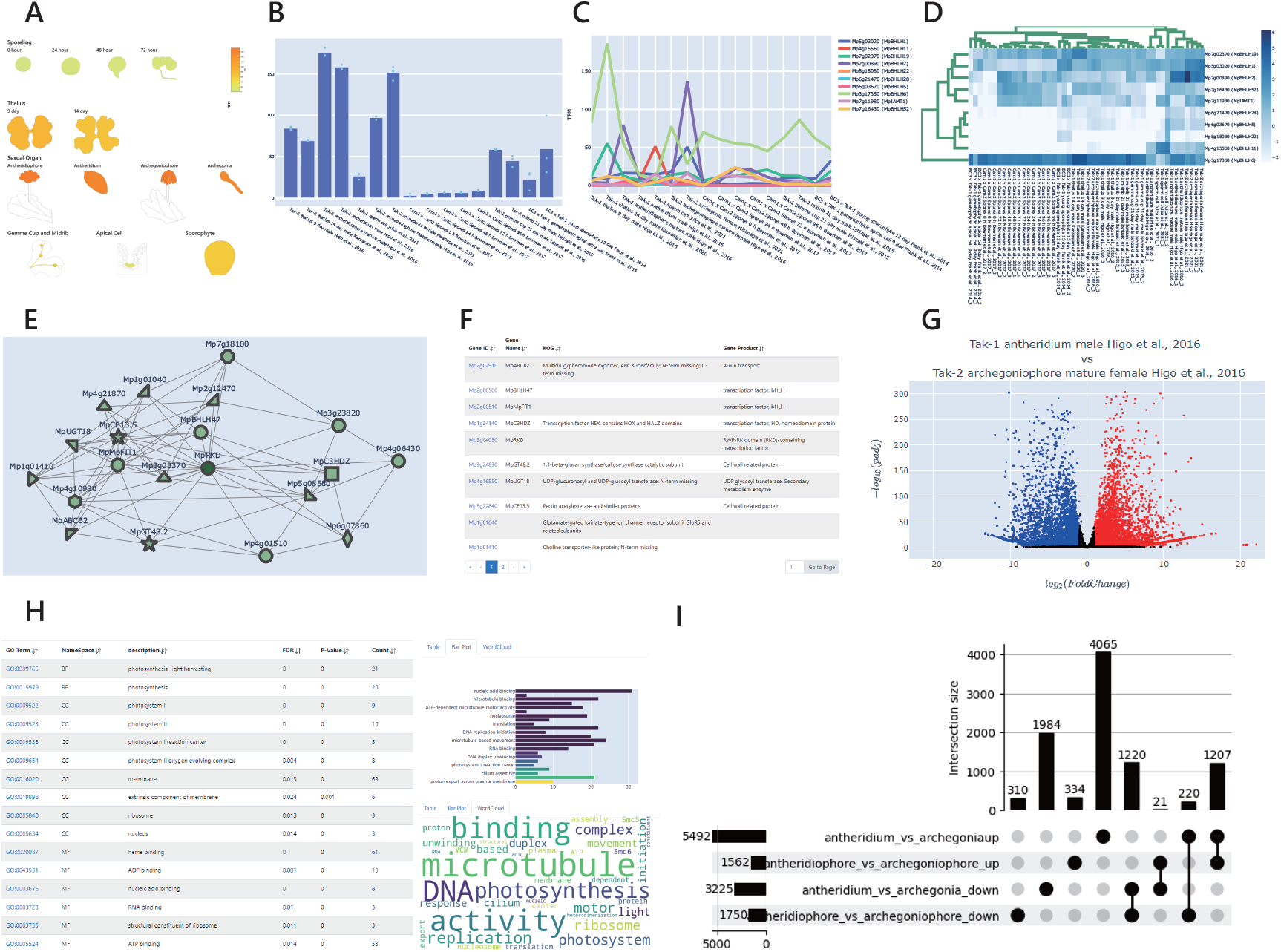
Basic tools in MBEX. (**A**) ‘Chromatic Expression Images’ shows the expression levels of a given gene in various tissues/organs. (**B**) ‘Bar Plot’ of expression levels of a given gene under a selected set of conditions. (**C**) ‘Line Plot’ of expression levels of a given set of genes under selected conditions. (**D**) ‘Clustergram’ of a given set of genes under a selected set of conditions. Expression levels are given in TPM, which can be downloaded in the CSV format. ‘Bar Plot’, ‘Line Plot’, and ‘Clustergram’ can be saved in the SVG, PNG, or JPEG formats. (**E**) ‘Co-expression Network’ generates a network of genes co-expressed with a selected gene. (**F**) A list of co-expressed genes with rank information and simplified annotation can be downloaded in the CSV format. (**G**) ‘Differential Expression’ generates a volcano plot showing genes differentially expressed in a pair of published datasets, where red, blue, and black points represent upregulated, downregulated, and non-differentially expressed genes, respectively. (**H**) ‘Enrichment’ provides a table, bar plot, and word cloud of enriched terms for a given set of genes. The list of enriched terms can be downloaded in CSV format. (**I**) ‘Set Relation’ generates an UpSet plot to show the numbers of genes in intersecting sets.

### Co-expression Tools

We provide two types of tools for co-expression analysis: visualization of co-expression networks and summarization of co-expression data in a tabular format (Fig. 3E, F).

‘Co-expression Network’ draws a co-expression network of a gene of interest with its co-expressed neighbors (within a distance of 3) genes. In contrast, ‘Functional Network’ lets users specify multiple genes as inputs to examine their connections within a co-expression network that is more expansive (up to a distance of 10) than that generated by ‘Co-expression Network’, and also filters the neighbor genes by annotations with words of interest. ‘NetworkDrawer’ is useful to depict a co-expression network represented by genes specified by users. Users can also specify the color for each marker in the diagram created by ‘NetworkDrawer’. In these diagrams, the nodes in the network appear in different shapes, which represent the functional category of the assigned KOG. The annotation information can be displayed when placing the mouse pointer on a shape. Details of the genes shown in the network are also available as a list, which includes MpGene ID, nomenclature information, descriptions of KOG, KEGG, and Pfam, the distance from the user-selected gene, the link to ‘OrthoPhyloViewer’, and the external link to its page in MarpolBase.

Other tools, ‘Co-Expression Table’ and ‘Rank Table’, are also available to display co-expression data in a tabular format. ‘Co-Expression Table’ presents the Pearson correlation coefficients and p-values of genes co-expressed with a user-specified gene under user-specified conditions. ‘Rank Table’ identifies genes whose transcription is correlated in all experimental conditions in relation to a given gene and displays their HRR, MR, and PCC with their MpGene ID, nomenclature information, descriptions of KOG, KEGG, and Pfam, the distance from the user-selected gene, and links to ‘Orthophyloviewer’ and MarpolBase.

### Analysis tools

Functional enrichment analysis using annotations such as GO, KEGG, KOG, and Pfam is often helpful to interpret the biological properties of a set of genes, such as co-expressed or differentially expressed genes in specific conditions. There are web-based tools for some model species to perform functional enrichment analysis by providing only the IDs or names of genes of interest (Mi et al., 2021). MBEX allows users to perform functional enrichment analysis by providing a list of gene names or MpGene IDs of interest. This tool allows users to view the results containing functional annotations, p-values, and false discovery rates (FDR), a bar plot representing count and FDR, and a word cloud representing word frequency in significantly enriched annotations.

In RNA-seq analyses, the identification of differentially expressed genes (DEGs) is one of the most widely used analytic methods for understanding molecular mechanisms underlying specific biological processes. However, it is often difficult for molecular biologists to access and analyze large datasets from public data. ‘Differential Expression’ in MBEX lets users identify DEGs from pairwise combinations of all experimental conditions by an R package, DESeq2 (Love et al., 2014). This tool enables users to generate and view an interactive volcano plot (Fig. 3G), and retrieve the MpGene ID, fold change value (FC; base-2 logarithm converted), p-adjustment value, and a link to the MarpolBase page of any gene of interest. The corresponding MpGene ID can be displayed by hovering the mouse pointer over the dot. The analysis results, including the list of up- and down-regulated genes, and whole DEGs, can be downloaded. In addition, users can perform GO enrichment analysis (GOEA) using the result of DEG analysis directly from the ‘GO analysis’ button on the result page.

We also provide the ‘Set Relation’ tool to integrate and compare multiple results such as DEG or coexpression analyses. This tool can help users understand the intersections and unions of multiple sets of genes, such as a set of similarly up- or down-regulated genes in different samples. It takes two or more files containing MpGene IDs as the input and generates an UpSet plot and tables representing set relations with detailed gene information. GOEA can also be performed using genes included in the user-selected sets.

### Case Study: an oil body-specific gene network

In order to test the power of MBEX to gain biological insights, we focused on genes associated with oil bodies in *M. polymorpha.* Oil bodies in liverworts, which are distinct from ‘oil bodies’ in angiosperms, are a synapomorphic feature of liverworts (Romani et al., 2022). In *M. polymorpha,* oil body cells are scattered in the plant body (Fig. 4A) and accumulate secondary metabolites such as terpenoids and bisbibenzyls, which serve as deterrents against herbivores (Romani et al., 2020; Kanazawa et al., 2020). Some terpene synthases are expressed in oil body cells in *M. polymorpha* (Suire et al., 2000; Takizawa et al., 2021), suggesting that they serve as factories and storage depots specialized for terpenoids and their derivatives. Therefore, this particular cell type serves as a suitable subject for co-expression and related analyses and for evaluation of the tools we have developed. Furthermore, while some biosynthetic pathways and enzymes associated with the rich diversity of terpenoids and bisbibenzyls in liverworts have been identified, a large fraction remain unknown (Asakawa and Ludwiczuk, 2018). Elucidating the gene networks associated with this specialized type of cell should help us further understand the biosynthetic pathways and their regulatory mechanisms. Previous work demonstrated that the number of oil body cells in *M. polymorpha* parenchyma changes in response to environmental conditions (Romani et al., 2020; Tanaka et al., 2016). Recently, two transcription factors, *MpC1HDZ* and *MpERF13,* were identified as positive regulators of oil body cell differentiation (Kanazawa et al., 2020; Romani et al., 2020). In both cases, loss-of-function mutants lack oil body cells; furthermore, in the case of *MpERF13,* gain-of-function alleles increased oil body cell numbers. Therefore, oil body cells represent a tightly regulated subpopulation of cells in the *M. polymorpha* thallus. Co-expression and related analyses (Fig. 4B, Supplementary Text) using the RNA-seq data obtained from these samples should reveal a suite of genes associated with the oil body formation and metabolism.

**Fig. 4.**
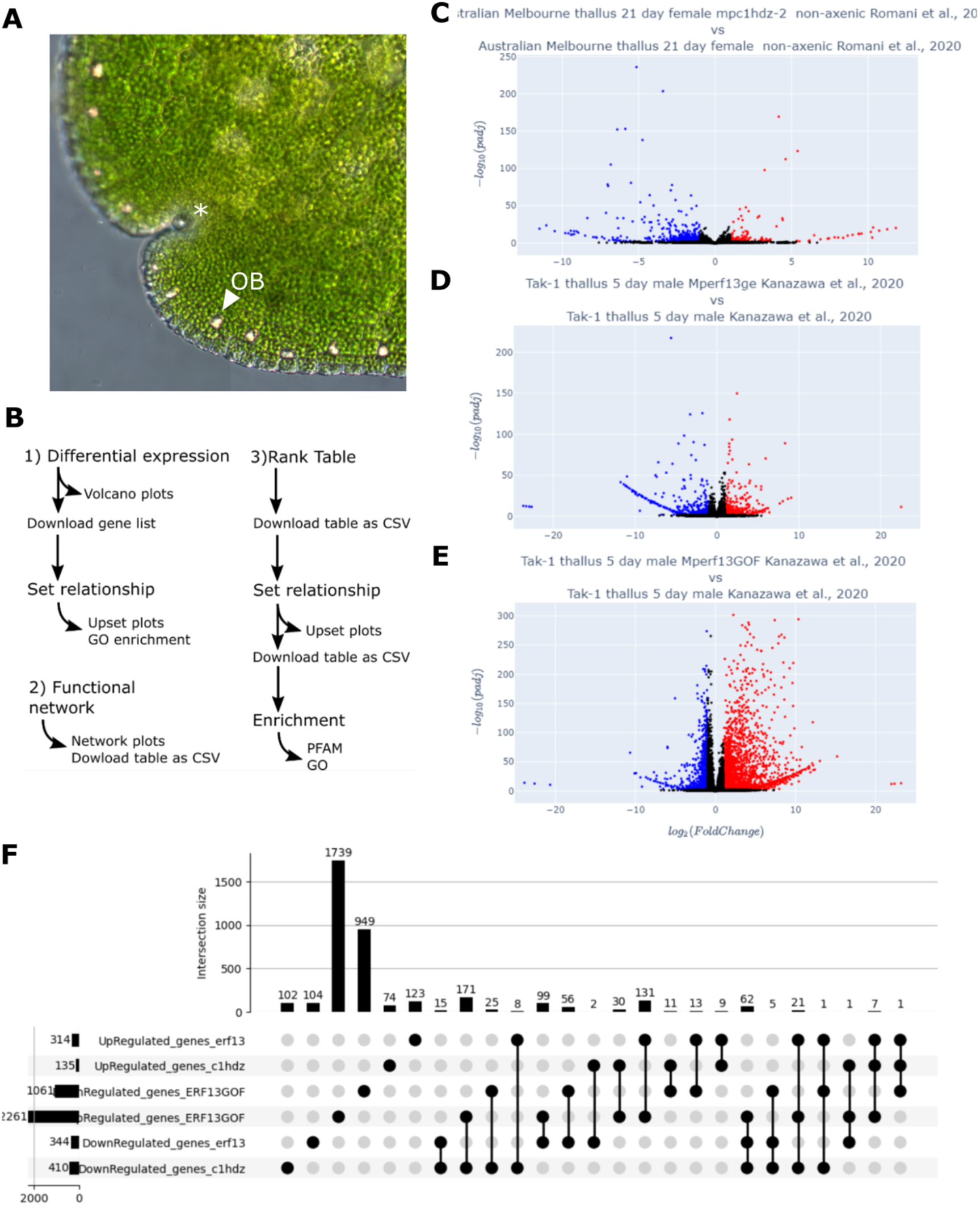
Flow of the case study on the oil body formation. (**A**) Oil bodies (indicated by a triangle) in a gemma. (**B**) Schematic overview of this analysis. (**C-E**) Volcano plots for comparisons between WT and loss-of-function mutants of *MpC1HDZ* and *MpERF13* (C and D, respectively), and between WT and gain-of-function mutants of Mp*ERF13* (E). (**F**) UpSet plot of up- and down-regulated genes in each mutant.

We reanalyzed RNA-seq experiments performed for loss-of-function *MpC1HDZ* mutants (Romani et al., 2020) and both gain-of-function (GOF) and loss-of-function *MpERF13* mutants (Kanazawa et al., 2020). First, to identify DEGs and their presumed functional categories, the corresponding RNA-seq datasets were selected and analyzed with the ‘Differential Expression’ tool, which generates volcano plots (Fig. 4C-E). The lists of DEGs were provided to the ‘Set Relation’ tool to generate an UpSet plot that visualizes all the possible comparisons between down- and up-regulated genes (Fig. 4F). The most prominent changes in gene expression were observed in *Mperf13^GOF^*, which shows growth defects due to the overproduction of oil body cells (Kanazawa et al., 2020). The upregulated genes in *Mperf13^GOF^* consistently overlap with genes downregulated in the loss-of-function *Mpc1hdz* (171 genes) and *Mperf13* (99 genes) mutants, 62 of which were consistently down-regulated in both loss-of-function mutants. In summary, the ‘Differential Expression’ tool is quite competent to re-mine published RNA-seq data to obtain new insights.

We exploited the ‘Co-expression Tools’ to gain more holistic and robust insights. Given that oil bodies are more abundant in certain tissues and under certain growing conditions (Tanaka et al., 2016), integrating multiple RNA-seq experiments in co-expression networks could help capture expression profiles characteristic of particular cell types or growth conditions. Furthermore, the co-expression approach could identify genes that are masked in differential expression analysis where only a limited number of RNA-seq datasets are examined. Therefore, we followed a similar approach in co-expression analysis that was implemented previously to unravel tissue-specific expression programs in *Sorghum* (Turco et al., 2017). In addition to *MpERF13* and *MpC1HDZ,* we selected two other proteins known to be specifically expressed in oil body cells (Mp*ABCG1*, and *MpSYP12B)* as baits to identify other genes with a similar expression profile (Kanazawa et al., 2020, 2016; Romani et al., 2020).

The ‘Functional Network’ tool identified 25 genes with the four bait genes (Mutual Rank < 2000) that encode enzymes at different steps of ‘terpenoid biosynthesis’ as assessed by the word-filtering function and are candidates for oil body-specific enzymes. Both the cytosolic (mevalonate, MVA) and plastid (methylerythritol 4-phosphate, MEP) pathways are involved in the biosynthesis of isoprene (IPP) building blocks for terpenoids in liverworts (Fig. 5A) (Adam et al., 1998). Interestingly, all the genes involved in the MVA pathway were highly co-expressed with the bait genes. According to Adam et al. (1998), the MVA pathway is the preferred source of isoprene for sesquiterpene biosynthesis, while the MEP pathway supplies substrates for mono- and diterpene synthesis. Sesquiterpenes are specifically located in oil bodies in *M. polymorpha* (Tanaka et al., 2016), suggesting the MVA pathway should be active in oil body cells. The *Mpc1hdz* mutants, which lack oil bodies, are depleted of sesquiterpenes and the monoterpene limonene (Romani et al., 2020). *MpMTPSL2* and Mp*CPT* encode a microbial-type terpene synthase-like enzyme and *cis*-prenyltransferase in the limonene biosynthesis pathway, respectively (Kumar et al., 2016), and are highly co-expressed with the other oil body marker genes (Fig. 5B), indicating that monoterpenes are also synthesized and accumulate in oil bodies. Other terpene synthases are also highly co-expressed with the oil body marker genes, including Mp*FTPSL2* and Mp*FTPSL3,* which encode fungal-type terpene synthase-like enzymes involved in the biosynthesis of sesquiterpenes (Takizawa et al., 2021). The *MpFTPSL2* promoter is specific to oil bodies (Takizawa et al., 2021), showing some fungal-type terpene synthase genes can also be oil body-specific markers. Although plant-type terpene synthases (TPSs) function in diterpene and other terpene biosynthesis reactions and are expressed broadly in various plant tissues (Kumar et al. 2016), Mp*TPS3* and *MpTPS7* exhibited co-expression with the oil body markers, suggesting that they may also participate in the biosynthesis of oil body-specific compounds.

**Fig. 5.**
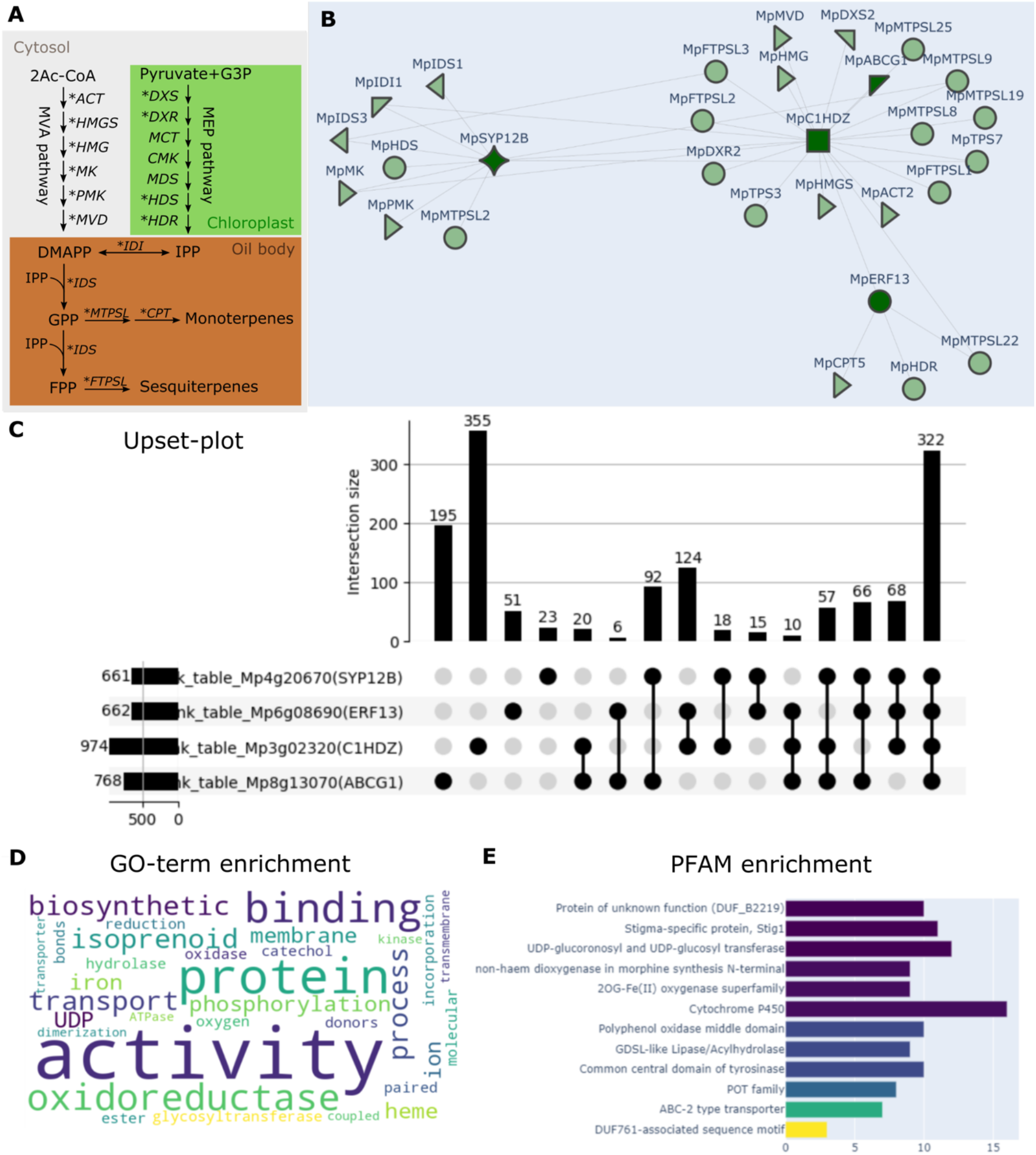
Functional network of the oil body genes and related enzyme genes. (**A**) The biosynthetic pathways of terpenoids. Different colors indicate the subcellular location of each pathway. Asterisks indicate genes represented in the functional network (B). ACT, acetoacetyl-CoA thiolase; CMK, CDP-ME kinase; CPT, cis-prenyltransferase; DMAPP, dimethylallyl diphosphate; DXR, 1-deoxy-d-xylulose 5-phosphate reductoisomerase; FPP, farnesyl diphosphate; DXS: 1-deoxy-D-xylulose-5-phosphate synthase; FTPSL, fungal-type terpene synthase-like; GPP, geranyl diphosphate; HDR, (*E*)-4-hydroxy-3-methylbut-2-enyl diphosphate reductase; HDS, (*E*)-4-hydroxy-3-methylbut-2-enyl diphosphate synthase; HMGR, 3-hydroxy-3-methylglutaryl-CoA reductase; HMGS, 3-hydroxy-3-methylglutaryl-CoA synthase; IDI, isopentenyl diphosphate isomerase; IDS, isoprenyl diphosphate synthases; IPP, isopentenyl diphosphate; MCT, 2-*C*-methyl-d-erythritol 4-phosphate cytidylyltransferase; MDS, 2-*C*-methyl-d-erythritol 2,4-cyclodiphosphate synthase; MEP, 2-*C*-methyl-d-erythritol 4-phosphate; MVD, mevalonate diphosphate decarboxylase; MK, mevalonate kinase; MTPSL, microbial-type synthase-like; PMK, phosphomevalonate kinase; TPS, terpene synthase. (**B**) The functional network of the four marker genes for oil bodies (dark green) and their functionally and transcriptionally associated genes (light green). The shape of each gene represents the KOG annotation, as explained on the MBEX website. (**C**) The UpSet plot of co-expressed genes with the four marker genes. (**D**) Word cloud of the GO enrichment result of genes, shown in arbitrary colors, that were coexpressed with at least three of the four marker genes. The size of each word indicates the degree of enrichment. (**E**) Bar plot of Pfam enrichment genes co-expressed with at least three of the four marker genes.

To explore other genes co-expressed with the oil body markers, a list of top co-expressed genes for each marker (Mutual Rank < 1000) was created by the ‘Rank Table’ tool, and the ‘Set Relations’ tool was used to identify genes that are co-expressed with more than one oil body marker. A large proportion of genes that co-expressed with each marker were also co-expressed with other markers, with 322 genes coexpressed with all four markers, confirming their similar expression patterns (Fig. 5C). In addition, this gene set also showed a consistent overlap with the genes identified in the ‘Differential Expression’ analysis.

A set of genes that co-expressed with at least three of the four baits was selected to inspect the functional aspects of the genes of inferred association with the oil body program. The ‘Enrichment’ tool for GO and Pfam enrichment analysis showed that the most frequent GO terms for biological processes were consistently associated with oil body-specific metabolism, including oxidoreductase, catechol oxidase, UDP-glycosyltransferase activities, and isoprenoid biosynthesis (Fig. 5D). At the same time, protein families including cytochrome P450, UDP glucuronosyl, UDP-glucosyl transferase, and ABC-2 type transporters were also enriched, as expected (Fig. 5E).

In addition to terpenoids, oil bodies also accumulate bisbibenzyl compounds, such as marchantin A and perrottetinene, which are unique to liverworts and are of economic interest (Asakawa and Ludwiczuk, 2018; Gülck and Møller, 2020). Their biochemistry is related to the phenylpropanoid pathway, but the enzymes involved are unknown. Several genes involved in phenylpropanoid biosynthesis were identified among the oil body co-expressed genes. Since a cytochrome P450 enzyme plays a critical role in bisbibenzyl biosynthesis (Friederich et al., 1999), cytochrome P450 genes within the functional network of oil bodies are good candidates for those responsible. In addition, genes encoding cytochrome P450s and UDP-glycosyltransferases that are co-expressed with the oil body markers could also contribute to the tailoring steps in terpenoid biosynthesis. It should be noted that the ‘Functional Network’ can also identify transcription factors involved in a given network. The ‘Functional Network’ analysis (Mutual Rank < 30) with the oil body marker genes, *MpERF13, MpC1HDZ, MpABCG1,* and *MpSYP12B,* revealed that an R2R3-MYB transcription factor gene, *MpMYB02,* is closely associated with *MpSYP12B,* suggesting its involvement in oil body functions. Indeed, *MpMYB02* was shown experimentally to be involved in bisbibenzyl biosynthesis (Kubo et al., 2018), and it is specifically expressed in oil body cells (Kanazawa et al., 2020).

Overall, this series of analyses support the idea that oil bodies work as cellular factories of secondary metabolites, and we successfully identified key genes likely to be part of the oil body-specific program. Both differential expression and co-expression approaches complement each other to strengthen predictions and suggest appropriate candidate genes. We also provide a step-by-step guide for reproducing this analysis on the MBEX website (Supplementary text). The workflow presented here can be easily adapted to investigate transcription programs associated with other cell types and metabolic pathways in *M. polymorpha*.

### Conclusion and Future Remarks

MBEX allows users to perform a series of analyses from fundamental data processing to data visualization on the Web. We anticipate that MBEX will evolve into a comprehensive and all-in-one analytical platform by continually incorporating newly published RNA-seq data and annotations. It should accelerate molecular biological discoveries in the liverwort *M. polymorpha* and place them in the context of land plant evolution.

## Materials and Methods

### Database Construction

Datasets used in MBEX were collected from SRA, and selected using the search condition of taxonomy ID *‘3169(=Marchantia polymorpha)’* and Library Strategy ‘RNA-seq’. SRA files were downloaded and converted into fastq files by using fasterq-dump (Sequence Read Archive Handbook) using default parameters. Quality control and trim of low-quality reads and adaptors were performed with fastp using default parameters. Trimmed reads were pseudo-aligned to the predicted transcripts from the representative gene models of the *M. polymorpha* Tak v6 genome using Salmon v1.4.0 (Patro et al., 2017) with the parameters ‘-1 A –validateMapping –seqBias –gcBias’. RNA-Seq counts were converted into non-normalized raw counts and Transcript Per Million (TPM) values per gene using the tximport R package. We implemented the backend and crawler of MarpolBase Expression in Python with the Django and FastAPI web framework. The data are stored in MySQL, PostgreSQL, MongoDB, and Redis databases. The frontend was developed in the TypeScript with React and Bootstrap framework. The plot’s dynamic and interactive elements are drawn using SVG markup language and TypeScript with the plotlyjs library.

### Correlation analysis

After log2 transformation with added 0.25, Pearson correlation coefficients (PCC) and p-values were calculated. Co-expression networks in MarpolBase Expression are based on Highest Reciprocal Rank (HRR) and Mutual Rank (MR). HRR for genes A and B is calculated as max(rank(PCC(A, B)), rank(PCC(B, A))), with 0 corresponding to the gene rank against itself. MR for genes A and B is calculated as √PCC(A, B) × PCC(B,A).

### Analysis of Differentially Expressed Genes

Differentially expressed genes were analyzed with the DESeq2 R packages (Love et al., 2014). First, a data frame was generated, including the expected non-normalized raw counts of only the samples present in the two groups to be compared and containing ≥ 3 biological replicates was performed using the estimateSizeFactors function, and the dispersion was calculated using the estimateDispersions function. nbionmWaldTest function was used to calculate differentially expressed genes.

### GO Enrichment and Functional Enrichment Analyses

Functional annotations, including GO terms, Pfam domains, and KEGG/KOG numbers were imported from MarpolBase. GOATOOLS (Klopfenstein et al., 2018) and in-house Python script with the SciPy library were used in GO Enrichment and Functional Enrichment analyses to detect over- and underrepresented terms based on Fisher’s exact test. The p-value was corrected by FDR using the Benjamini/Hochberg procedure.

### Orthogroup Clustering and Construction of Phylogenetic Trees

OrthoFinder v2.4.0 (Emms and Kelly, 2019) was used to group genes into orthogroups and construct phylogenetic trees, using Diamond to determine sequence similarities with default parameters.

## Funding

This research was funded by JSPS/MEXT KAKENHI [grant numbers: 17H07424, 19H05675, and 22H00417 to T.K., 20K15783 to Y.T., and 16H06279 (PAGS) to T.K. and Y.N.].

S.K. was supported by a Grant-in-Aid for JSPS Research Fellows [21J15550]. J.L.B. was supported by Australian Research Council [DP200100225]. F.R. was supported by BBSRC [BB/T007117/1].

## Acknowledgments

The authors thank Miyuki Iwasaki for providing pictures used in the ‘Chromatic Expression Images’ tool. Computations were performed on the supercomputer of ACCMS at Kyoto University.

## Disclosures

All authors declare no conflict of interest regarding the contents of this article.

## Supplementary Materials

### Supplementary Text

#### OrthoPhylo Viewer

Users can visualize an orthologous group (orthogroup) of genes and their phylogenetic tree (Supplementary Fig. 1A, B). Five plant species are covered, i.e., *Arabidopsis thaliana, Oryza sativa, Physcomitrium patens, Marchantia polymorpha,* and *Klebsormidium nitens.* This link is available in the table view in the co-expression analysis.

#### DataSource

Users can search manually curated condition information and SRA samples used in MBEX. These data can be downloaded in CSV format. The raw count, TPM and correlation matrix are also available for download from the DataSource page.

#### Co-Ex Viewer

Users can visualize expression correlations between two genes (Supplementary Fig. 1D). Using the Co-Ex Viewer tool, users can examine whether two genes of interest are transcriptionally correlated or not under most or certain conditions. Furthermore, users can view the conditions under which the two selected gene pairs are co-expressed. By placing the pointer on the dot, users can see which condition the dot is in. This tool informs users about which conditions are outliers, and under which specific conditions, excluding outliers, the genes are co-expressed.

#### Step-by-step instructions to reproduce the Case Study

DEG Analysis

(“Analysis Tools” -> “Differential Expression”, or https://marchantia.info/mbex/diffexp)

1. Select and compare the following pair of conditions to generate a volcano plot and lists of DEGs

- *Australian Melbourne thallus 21 day female mpc1hdz-2 non-axenic Romani et al., 2020* and *Australian Melbourne thallus 21 day female non-axenic Romani et al., 2020*
2. Download lists of both UpRegulated and DownRegulated gene IDs separately for subsequent analysis from “Download Gene List”
3. Repeat steps 1-2 above for the following pairs

- *Tak-1 thallus 5 day male Mperf13GOF Kanazawa et al., 2020* and *Tak-1 thallus 5 day male Kanazawa et al., 2020*
- *Tak-1 thallus 5 day male Mperf13ge Kanazawa et al., 2020* and *Tak-1 thallus 5 day male Kanazawa et al., 2020*

Set Relation Analysis for DEG analysis

(“Analysis Tools” -> “Set Relation”, or https://marchantia.info/mbex/setrel)

1. Upload all of the six files that were downloaded in the previous step
2. Click “Submit” to obtain DEGs co-expressed under the conditions selected above

Functional Network Tools for analyzing terpene biosynthesis (https://marchantia.info/mbex/functionalnetwork)

1. Set parameters under “Config” as follows

1. Rank Type: MR (Mutual Rank)
2. Rank Cutoff: 2000
2. Set gene names MpERF13 (Mp6g08690), MpC1HDZ (Mp3g02320), MpABCG1 (Mp8g13070), MpSYP12B (Mp4g20670)
3. Set filter word as terpene

Rank Table Analysis to explore genes co-expressed with oil body markers

(https://marchantia.info/mbex/ranktable)

1. Set parameters under “Config” as follows

1. Rank Type: MR (Mutual Rank)
2. Rank Cutoff: 1000
2. Set gene name MpERF13 and “Submit”
3. “Download Table as CSV” to save the result as a CSV file for the subsequent analysis.
4. Repeat steps 2-3 for MpC1HDZ, MpABCG1, MpSYP12B

Set Relation Analysis for genes co-expressed with oil body markers (https://marchantia.info/mbex/setrel)

1. Upload the 4 CSV files downloaded in the previous step and “Submit”
2. Check at least three sets of genes co-expressed
3. “Download Selected Gene Information” to save the result CSV file for the subsequent analysis

Functional Enrichment Analysis (https://marchantia.info/mbex/enrichment)

1. Upload the file downloaded in the previous step and “Submit”
2. Set Annotation Type as GO for GO Enrichment
3. Enter gene names of interest and “Set Gene Names”

## Supplementary Figures

**Supplementary Fig. S1.**
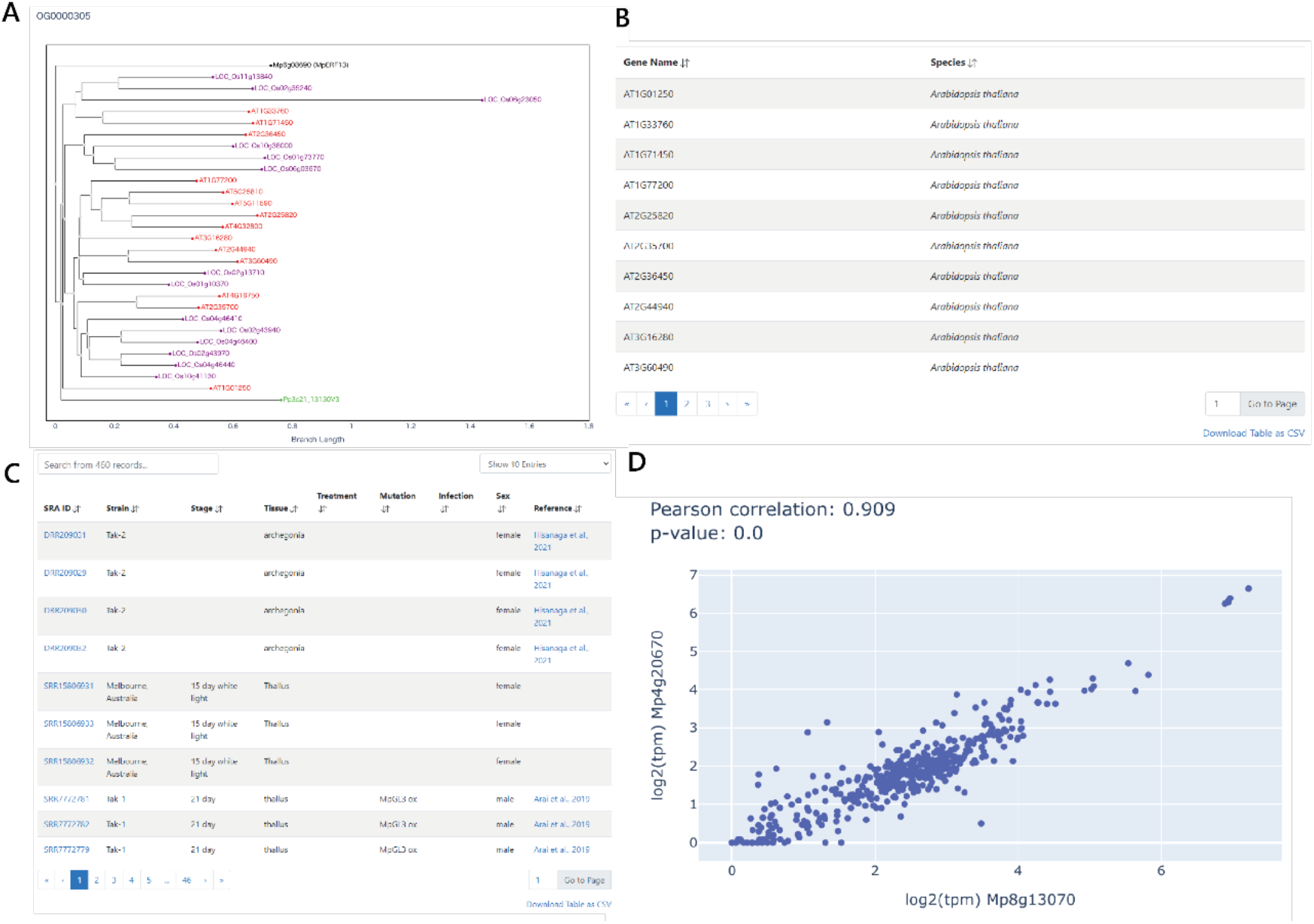
Extended tools in MBEX. (**A**) “OrthoPhyloViewer” shows a phylogenetic tree of the orthogroup for a given gene. (**B**) A list of genes in the orthogroup. (**C**) “Data Source” lists all the SRA samples used in MBEX for download in the CSV format. (**D**) “Co-Expression Viewer” shows expression correlations between two genes in all samples used in MBEX.

**Supplementary Fig. S2.**
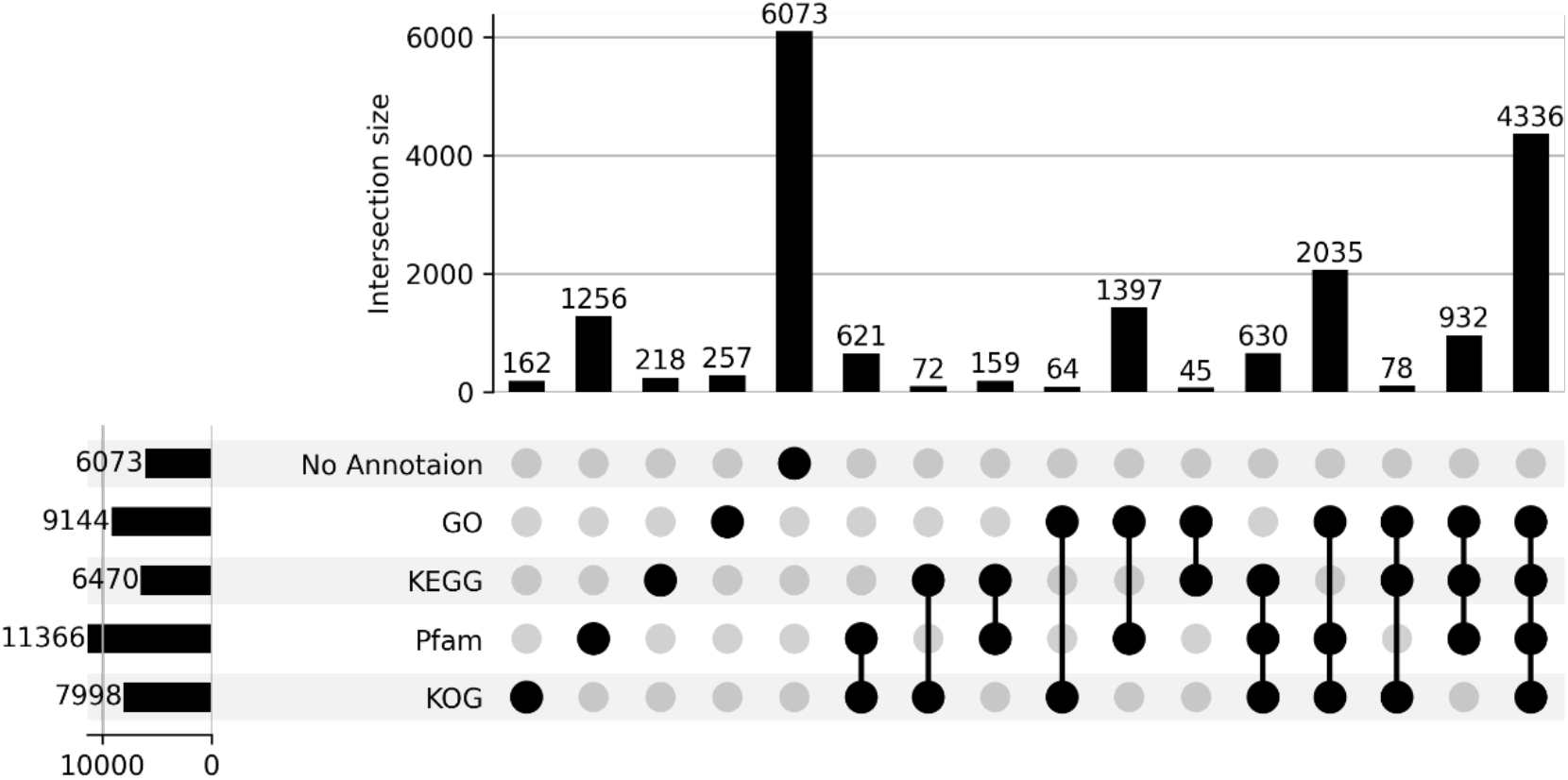
The annotation coverage of the *M. polymorpha* genes.

**Supplementary Fig. S3.**
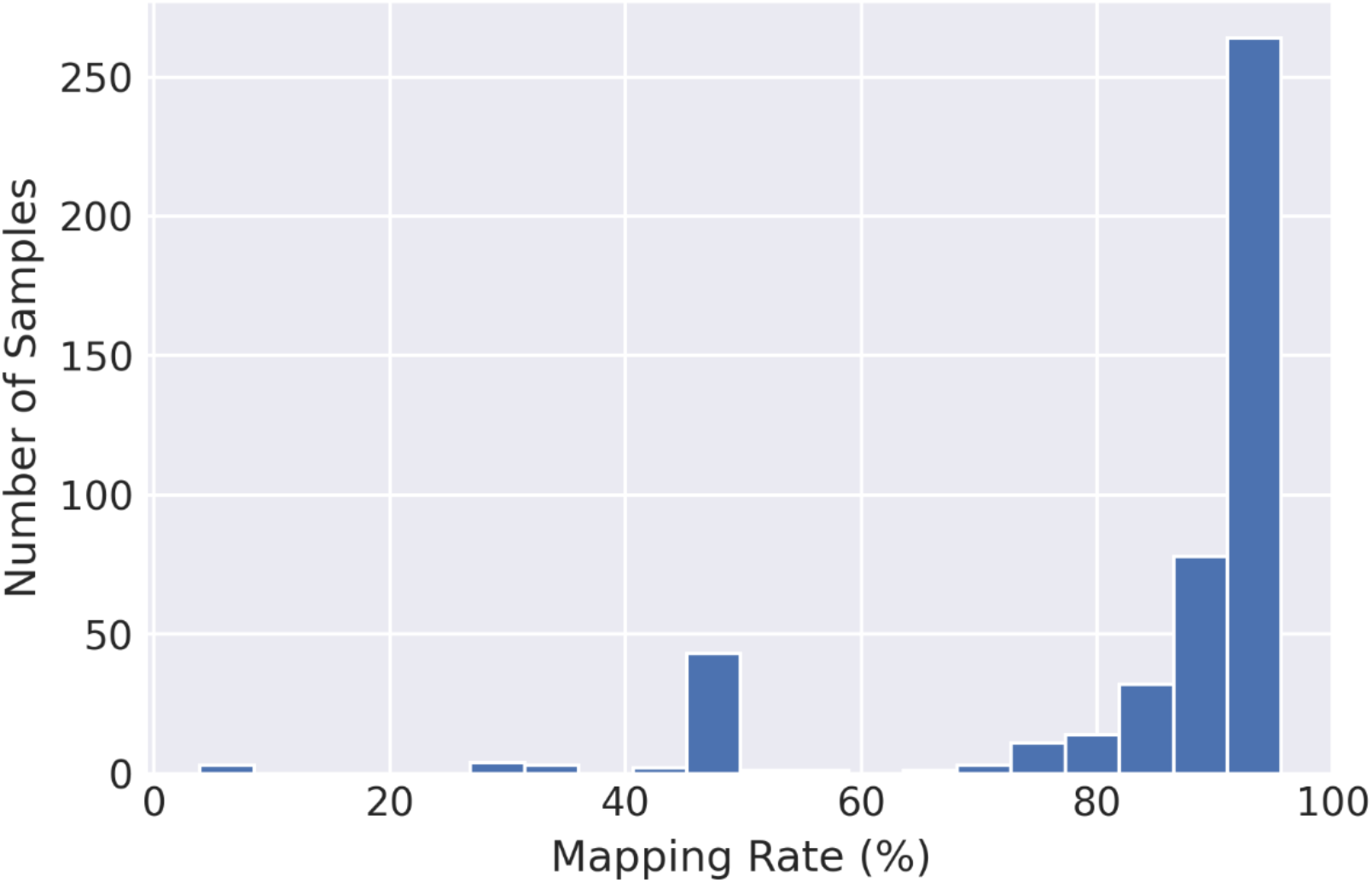
Distribution of the mapping rates for the RNA-seq data used in this study.

**Supplementary Fig. S4.**
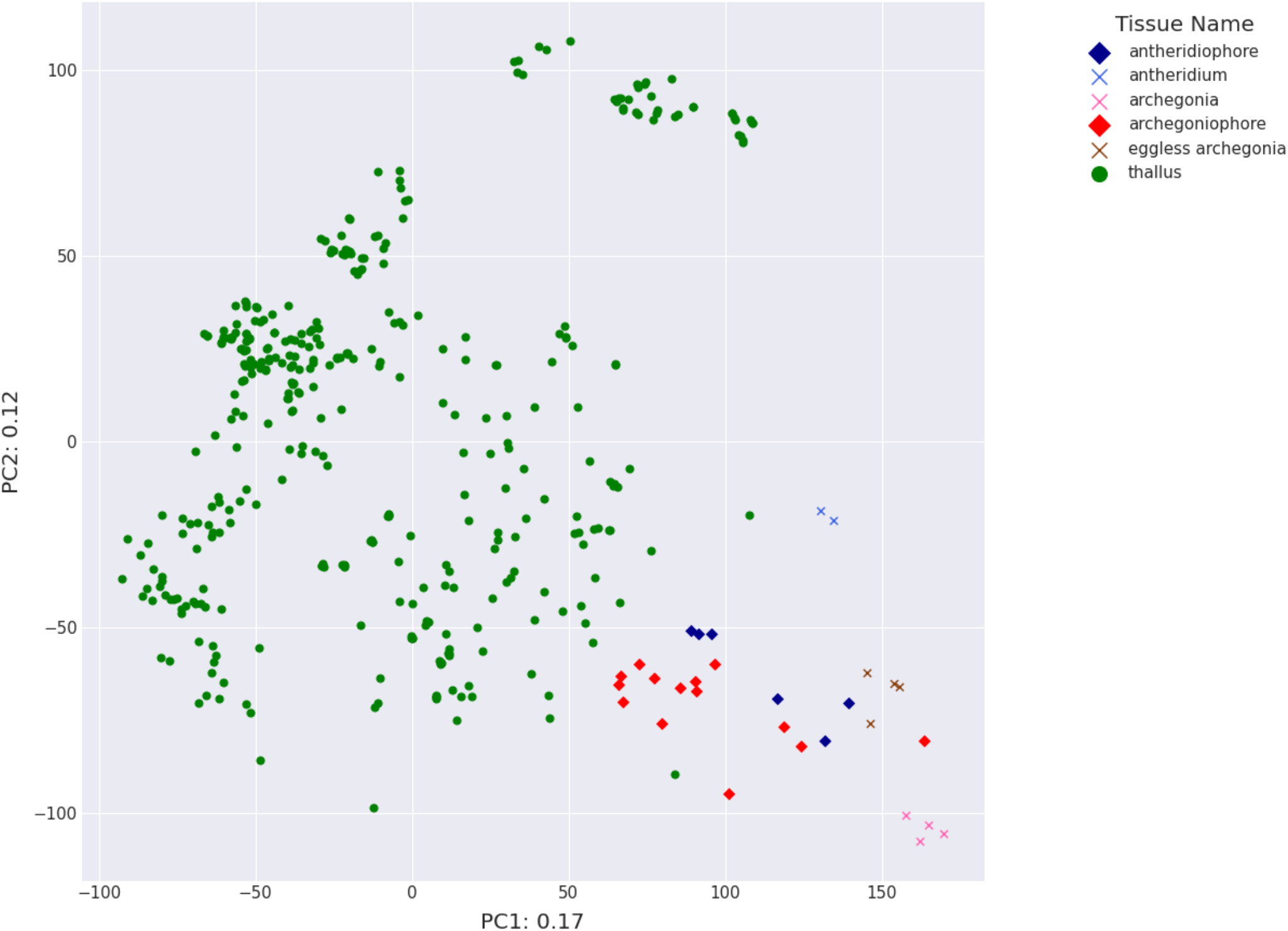
PCA analysis of thallus, gametangiophore, and gametangia.

## Supplementary Table

**Supplementary Table S1.** Sample number of each tissue.

| Tissue Name | Sample Number |
| --- | --- |
| antheridiophore | 6 |
| antheridium | 2 |
| archegonia | 4 |
| archegoniophore | 14 |
| eggless archegonia | 4 |
| gametophytic apical cell | 3 |
| gemma | 20 |
| gemma cup | 4 |
| midrib | 3 |
| mixed | 1 |
| sperm cell | 3 |
| sporelings | 13 |
| spores | 3 |
| sporophyte | 4 |
| thallus | 376 |

## Supplementary Data

**Supplementary Data** Mapping rates of the samples used in this study (in a separate CSV file).

## Notes

### Competing Interest Statement

The authors have declared no competing interest.

